# Switch costs in inhibitory control and voluntary behavior: A computational study of the antisaccade task

**DOI:** 10.1101/313643

**Authors:** Eduardo A. Aponte, Klaas E. Stephan, Jakob Heinzle

## Abstract

An integral aspect of human cognition is the ability to inhibit habitual responses in order to initiate complex, rule-guided actions. Moreover, humans have also the ability to alternate between different sets of rules or tasks, at the cost of degraded performance when compared to repeating the same task, a phenomenon called the ‘task switch cost’. While it is recognized that switching between tasks requires often to inhibit habitual responses, the interaction between these two forms of cognitive control has been much less studied than each of them separately. Here, we use a computational model to draw a bridge between inhibitory control and voluntary action generation and thereby provide a novel account of seemingly paradoxical findings in the task switch literature. We investigated task switching in the mixed antisaccade task, in which participants are cued to saccade either in the same or in the opposite direction to a peripheral stimulus. Our model demonstrates that stopping a habitual action leads to increased inhibitory control that persists on the next trial. However, enhanced inhibition affects only the probability of generating habitual responses, and, contrary to previous accounts, cannot be characterized as proactive task interference. In addition, our model demonstrates that voluntary actions (but not habitual responses) are slower and more prompt to errors on switch trials compared to repeat trials. We conclude that precisely the interaction between these two effects explains a variety of contradictory findings reported in the literature.

## Introduction

A hallmark of high-order cognition is the ability to alternate between different voluntary actions, as well as between habitual and non-habitual responses (Isoda and Hikosaka, 2008). However, alternating between different tasks engenders reaction time (RT) and error rate (ER) switch costs (Kiesel et al., 2010). While inhibitory control of habitual actions (Aron, 2011) and flexible action selection (Monsell, 2003) have been investigated in great detail, the interplay between them and its impact on task switching has received much less attention (but see Hikosaka and Isoda 2010). Saliently, while great effort has been devoted to developing computational models of action inhibition (Schall et al., 2017) and task switching (Karayanidis et al., 2010; Schmitz and Voss, 2014), models of the interaction between these two forms of cognitive control have been less prominent in the literature.

An attractive experimental paradigm to study the above phenomena is the antisaccade task (Hallett, 1978; Munoz and Everling, 2004), in which a habitual response – a prosaccade towards a salient peripheral stimulus – needs to be overwritten by a non-habitual action, i.e., an antisaccade in the opposite direction of the stimulus. Behaviorally, switch costs in the mixed antisaccade task, in which pro- and antisaccade trials are alternated, have been investigated in great detail (Barton et al., 2002; Cherkasova et al., 2002; Hunt and Klein, 2002; Manoach et al., 2002; Bojko et al., 2004; Fecteau et al., 2004; Manoach et al., 2004; Barton et al., 2006a; 2006b; Manoach et al., 2007; Rivaud-Pechoux et al., 2007; Ansari et al., 2008; Ethridge et al., 2009; Franke et al., 2009; Mueller et al., 2009; Lee et al., 2011; Weiler and Heath, 2012a; 2012b; DeSimone et al., 2014; Weiler and Heath, 2014a; 2014b; Heath et al., 2015; Pierce et al., 2015; Heath et al., 2016; Chan et al., 2017). Despite the large number of studies, no unified picture of the cost of switching in this paradigm has emerged. In particular, all human studies we are aware of have reported higher latencies in switch prosaccades (i.e., correct prosaccades that follow an antisaccade trial) than in repeat prosaccades. The costs associated with switch antisaccades are less clear: While some studies have indicated that switch antisaccades display lower RT than repeat trials (e.g., Cherkasova et al., 2002), others have reported both lower and higher RT (e.g., Barton et al., 2006a), and yet others indicate no switch costs (e.g., Weiler and Heath, 2012a).

From a theoretical perspective, two main explanations for switch costs in the antisaccade task have been proposed. According to the task-set reconfiguration hypothesis (Rogers and Monsell, 1995; Barton et al., 2006a), switch trials require the active reconfiguration of the task-set relevant to the new trial. This process is assumed to be an act of endogenous control that is not necessary in repeat trials, is time consuming, and can be prepared in advance of the peripheral stimulus. While intuitively appealing, this hypothesis is at odds with the observation that switch antisaccades are sometimes *faster* than repeat antisaccades (Cherkasova et al., 2002). By contrast, the task inertia hypothesis (Allport et al., 1994; Barton et al., 2006b; Weiler et al., 2015) postulates that passive interference caused by non-dominant rules (antisaccades) lead to pro- but not antisaccade RT switch costs. In other words, antisaccades require the activation of a ‘non-dominant’ rule, which interferes with prosaccades on the next trial. Because prosaccades are the ‘dominant’ rule, no interference occurs after this task-set has been activated. Again, this hypothesis is at odds with positive switch costs in switch pro- and antisaccades (Barton et al., 2006a). In other words, none of these hypotheses offers a satisfying explanation of the conflicting behavioral findings in the antisaccade task.

One approach to reconcile conceptual theories and seemingly contradictory experimental evidence is the application of generative models to empirical data (Monsell, 2003; Karayanidis et al., 2010; Heinzle et al., 2016), which might help disentangle the mechanisms behind switch costs. In this direction, we recently developed the *Stochastic Early Reaction, Inhibition and late Action* (SERIA) model (Aponte et al., 2017) of the antisaccade task. In essence, SERIA combines the ‘horse-race’ model of the countermanding saccade task (Logan et al., 1984; Camalier et al., 2007) to explain the inhibition of habitual, fast prosaccades, with a second race between two voluntary, or rule-guided actions that generate pro- and antisaccades. In contrast to previous models (Noorani and Carpenter, 2013), SERIA takes into account that prosaccades are not only reactive or habitual saccades, but can also be the result of a rule-guided decision process.

The main goal of our study was to investigate whether switch costs can be attributed to the inhibition of habitual responses and/or to the generation of voluntary saccades. Moreover, we investigated whether our modeling supports and explains the predictions of the task inertia and/or the task reconfiguration hypotheses. With these goals in mind, we applied SERIA to two versions of the antisaccade task (Aponte et al., 2018). In Task 1, the peripheral stimulus served simultaneously as task cue, indicating whether a pro- or an antisaccade should be performed. In Task 2, subjects were cued about the task demands in advance of the peripheral stimulus. Following previous reports, we expected positive antisaccade RT switch costs in Task 1, in which task and direction cue overlapped (similar to the short delay condition in Hunt and Klein, 2002; Barton et al., 2006a; Ethridge et al., 2009; see also Meiran, 1996). In Task 2, we expected either a negative or non-significant antisaccade switch cost, as the task cue was presented much in advance of the peripheral saccade target (Barton et al., 2006a; Ethridge et al., 2009; DeSimone et al., 2014).

Our results indicate that switch costs in the antisaccade task are explained by two distinct inter-trial effects that impact inhibitory control and voluntary action generation independently. Specifically, SERIA demonstrates task-inertia *like* effects on inhibitory control of habitual actions, as well as task-set reconfiguration costs in the execution of voluntary actions. We show here that by distinguishing between inhibitory control and voluntary action generation, it is possible to develop a unified account of the cost of switching in the antisaccade task that explains empirical findings and reconciles previous theoretical accounts.

## Methods

In this study, we analyzed switch costs in the data reported in Aponte et al. (2018), and hence we provide here only a short summary of the experimental procedures. The data is available for download at doi:10.3929/ethz-b-000296409. This experiment was approved by the ethics board of the Canton of Zurich, Switzerland (KEK-ZH-Nr.2014-0246) and was conducted according to the Declaration of Helsinki.

### Participants

Twenty-five healthy male subjects participated in the experiment. All subjects had normal or corrected to normal vision and provided written informed consent to participate in the study.

### Apparatus

The experiment was conducted in a dimly illuminated room. Subjects sat 60cm in front of a computer screen (41.4×30cm; Philips 20B40; refresh rate 85Hz). Eye position was recorded at a sampling rate of 1000Hz with a remote, infrared eye tracker (Eyelink 1000; SR Research, Ottawa, Canada). Head position was stabilized using a chin rest. The experiment was controlled by in-house software written in the Python programming language (2.7) using the PsychoPy package (1.82.02; Peirce, 2007; 2008).

### Experimental design

Subjects took part in two tasks consisting of three blocks of mixed pro- and antisaccade trials. Each block comprised 200 trials of which either 20, 50, or 80% were prosaccade trials. Before the main experiment, subjects underwent a training block of 50 prosaccade trials followed by 50 antisaccade trials for each task. During training (but not in the main experiment), subjects received feedback about their performance.

In Task 1 (Fig. 1), two red circles (radius 0.25°) were presented throughout the experiment at an eccentricity of ±12°. Each trial started with a central fixation cross (0.6×0.6°). Subjects were required to fixate for at least 500ms, after which a random interval of 500 to 1000ms started. Completed this period, the fixation cross disappeared, and a green bar (3.48×0.8°) in either horizontal or vertical orientation was presented centered on one of the red circles for 500ms. Subjects were instructed to saccade to the red circle cued by a horizontal green bar (prosaccade trials), and to saccade to the un-cued circle in case of a vertical bar (antisaccade trials). The next trial started after 1000ms. Pro- and antisaccade trials were randomly interleaved, but the same sequence was presented to all subjects. The location (left of right) of the peripheral cue was randomly permuted, such that the number of pro- and antisaccade trials in each direction was the same.

**Figure 1:**
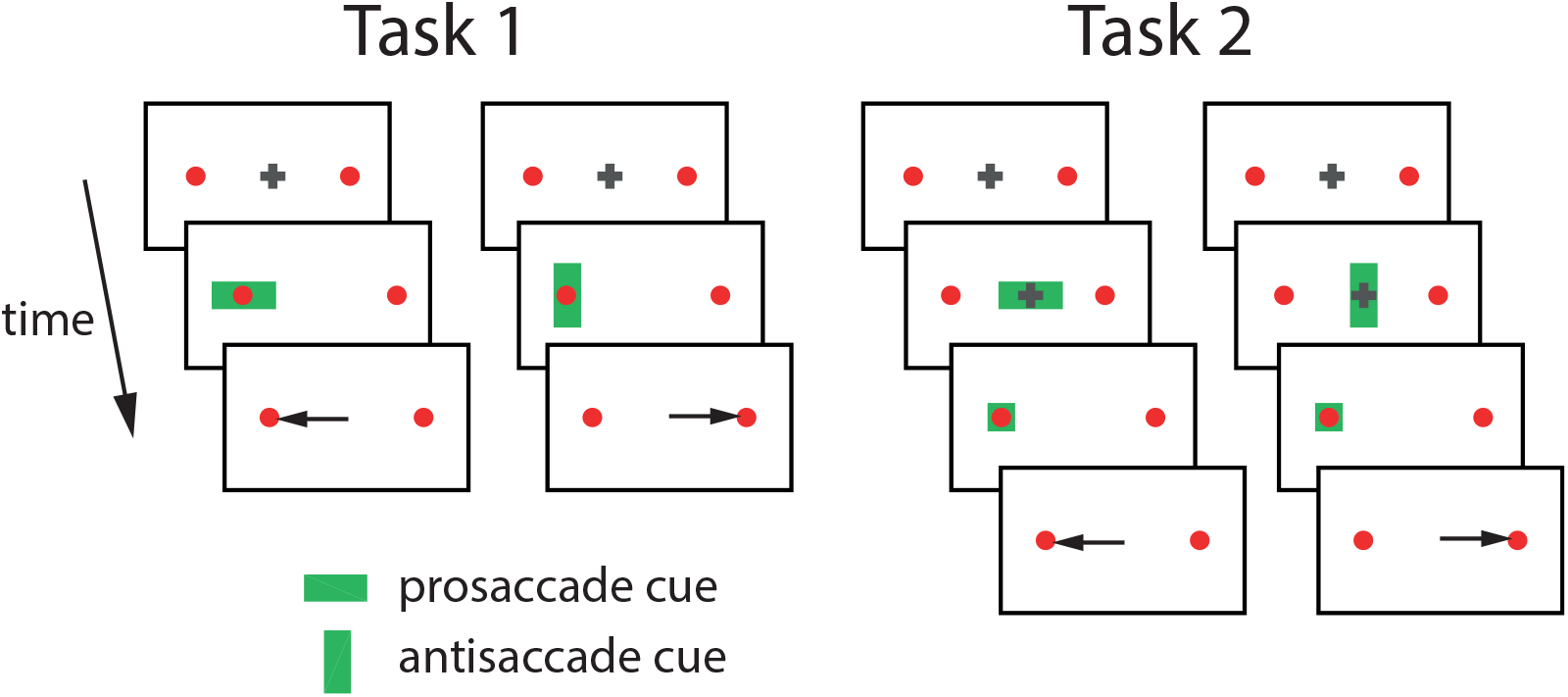
Experimental design. In both tasks, participants were instructed to first fixate to a central cross. **Task. 1:** After a variable interval (500-1000ms), a cue indicating the trial type was presented behind one of the peripheral red circles for 500ms. Depending on the cue orientation a saccade had to be performed toward or away from the cued target. **Task. 2:** Before the peripheral stimulus was presented, subjects were cued for 700ms about the task to be performed. After this cueing period, the central fixation cross disappeared, and a neutral cue was presented behind one of the peripheral red circles for 500ms. Depending on the orientation of the central green bar, a saccade toward or away from the cued target had to be performed.

Task 2 differed in that subjects were cued about the trial type in advance of the peripheral stimulus. As in Task 1, subjects were required to initially fixate a grey cross for 500 to 1000ms. After this interval, either a horizontal or a vertical bar was displayed behind the fixation cross. The bars had the same dimensions and color in both tasks. 700ms later, the green bar and the fixation cross were removed, and a green square (1.74°x1.74°) was presented centered behind one of the red circles for 500ms. Participants were instructed to saccade to the cued circle when a horizontal bar had been presented before and to saccade to the non-cued circle otherwise. The next trial started after 1000ms.

### Data processing

Saccades were detected with the software provided by the eye tracker manufacturer (Stampe, 1993), which uses a 22°/s and 3800°/s^2^ threshold to define the start of a saccade. Only saccades larger than 2° were included in the analysis. Trials were rejected in case of eye blinks or if subjects failed to maintain fixation before the peripheral cue was presented. Saccades with a latency above 800ms or below 50ms were rejected as invalid. Antisaccades were also rejected if their RT was less than 90ms. Only trials that directly followed a valid trial were included in the final analysis.

### Statistical Analysis

As variables of interest, we investigated mean RT of correct saccades and mean ER. These were analyzed with a generalized linear mixed effects (GLME) model implemented in the programming language *R* (package *lme4;* Bates et al., 2015). Independent variables were prosaccade trial probability (PP) with levels 20, 50 and 80%; trial type (TT); switch trial (SWITCH) with levels *switch* and *repeat*; and SUBJECT entered as a random effect. Significance was assessed through *F* tests with the Satterthwaite approximation to the degrees of freedom (Luke, 2017). For ER, the probit function was used as link function in the GLME. To test for significant effects, we used the Wald Chi-squared test implemented in the *car* package (Fox and Weisberg, 2011). When probabilities were investigated, we used a beta regression model implemented in the package *glmmADBM* (Fournier et al., 2012). Again, significance was tested with Wald Chi-squared tests.

### The SERIA model

Briefly, SERIA (Aponte et al., 2017) models the race of four independent accumulators or units: an early (*u*_e_), an inhibitory (*u_i_*), a late prosaccade (*u_p_*), and an antisaccade (*u*_a_) unit. An action *A* ∈ {*pro., anti.*} and its latency *T* ∈ [0, ∞[are treated as random variables, whose distribution is a function of the hit times of each of the units, *U_e_*, *U_i_*, *U_p_*, and *U_a_* respectively. Conceptually, SERIA can be decomposed into two different competitions (see Figure 2): First, the early unit, which models reactive, habitual responses, generates a prosaccade at time *t* if it hits threshold at time *t* (i.e., *U_e_* = *t*) before all other units. An early response can be stopped by the inhibitory unit if the latter hits threshold at some earlier point. In that case, either a pro- or an antisaccade is generated, depending on the outcome of the second race decision process between the late pro- and antisaccade units. For example, a late prosaccade at time *t* is generated if the late prosaccade unit hits threshold at *U_p_* = *t* before the antisaccade unit (i.e., *U_a_* > *t*).

**Figure 2:**
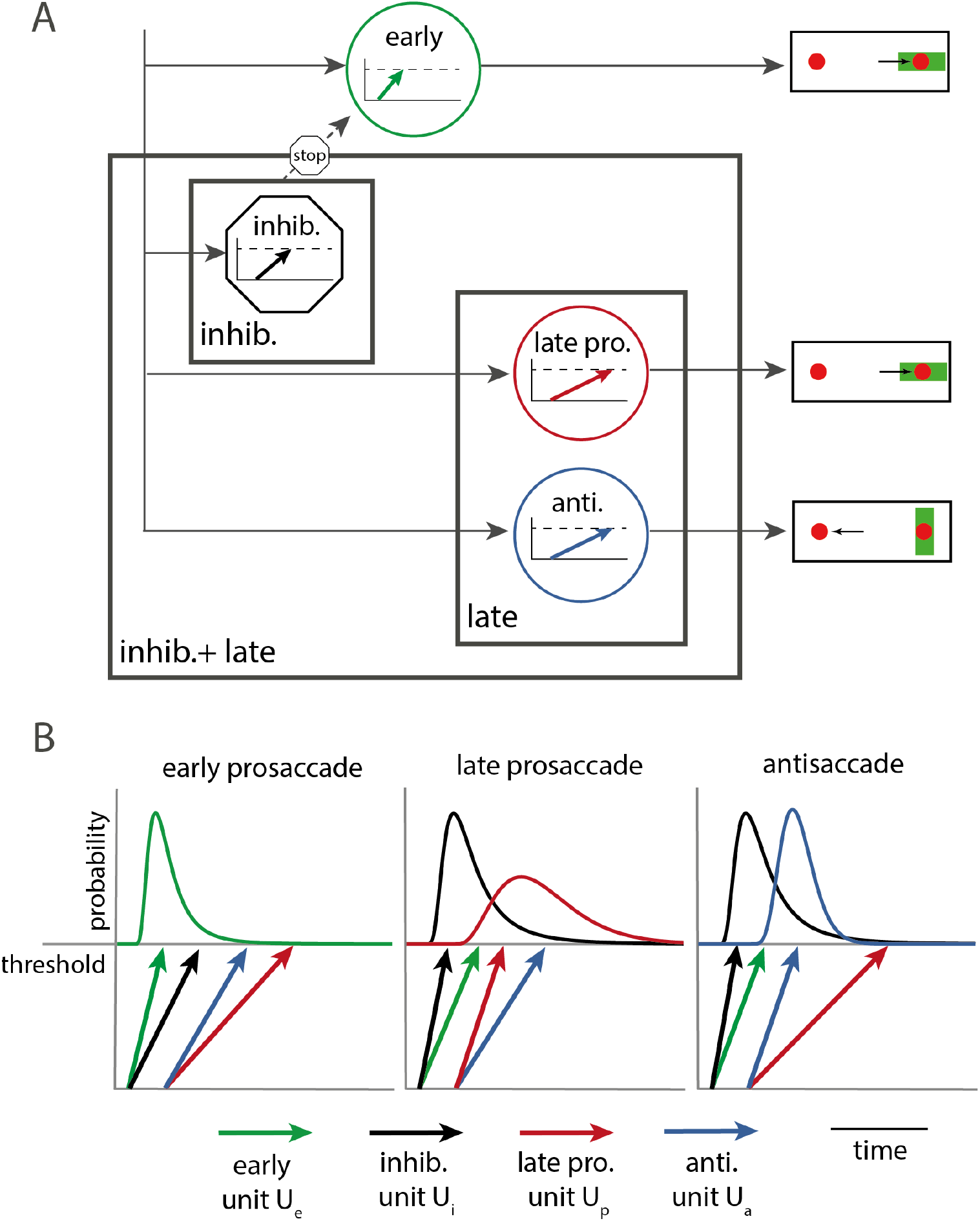
The SERIA model. **A.** SERIA is a race model that incorporates four different units (displayed as circles): an early prosaccade unit (green), an inhibitory unit (black), a late prosaccade (red) and an antisaccade unit (blue). We hypothesized that the effect of the previous trial could affect the inhibitory unit (*inhib.*), the late units (*late*), or both (*inhib.+late*). These three hypotheses are represented by black frames indicating the units affected by the previous trial under the corresponding hypothesis. **B.** The RT distributions are a function of the hit time distributions of the four units. Early reactions, which are always prosaccades, occur when the early unit hits threshold before all other units. A late prosaccade occurs mainly when the early unit is stopped by the inhibitory unit, and the late prosaccade unit hits threshold before the antisaccade unit. Similarly, antisaccades can only occur when the antisaccade unit hits threshold before the late prosaccade unit. Figure modified with permission from Aponte et al. (2018).

Concretely, SERIA provides an explicit formula for the probability of an action *A* and its RT. First, a prosaccade at time *t* is generated when either the early unit *u_e_* hits threshold at time *t* (i.e., *U_e_* = *t*) before all other units. The probability of this event is

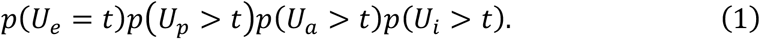

Furthermore, a prosaccade at time *t* can be triggered when the late prosaccade unit hits threshold at *t* before all other units

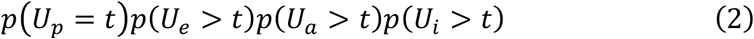

or when an early response is stopped by the inhibitory unit (i.e., *U_i_* < *t* and *U_i_* < *U_e_*), and the late prosaccade unit hits threshold before the antisaccade unit

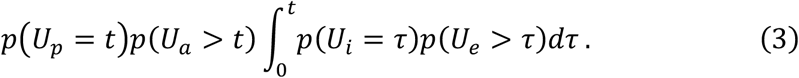

Similarly, an antisaccade at time *t* is generated when the antisaccade unit hits threshold at *t* (*U_a_* = *t*), before all other units

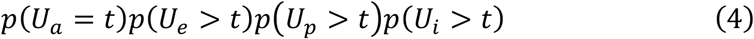

or when the antisaccade unit hits threshold before the late prosaccade unit after an early prosaccade has been stopped

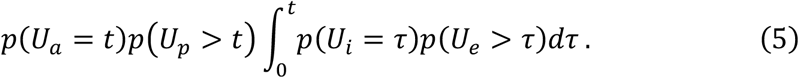

To fit the model, we assumed a parametric form for the hit times of each of the units: the hit times of the early (*U_e_*) and inhibitory unit (*U_i_*) were modeled with the inverse Gamma distribution, while the hit times of the late units (*U_p_* and *U_a_*) were modeled using the Gamma distribution (Aponte et al., 2017). Thus, each unit could be fully characterized by two parameters controlling the mean and variance of the hit times. Accordingly, 8 parameters were required for the 4 units in a given condition.

### Model space

We aimed to answer two different questions through quantitative Bayesian model comparison (Kass and Raftery, 1995; Stephan et al., 2009) and qualitative predictive fits (Gelman et al., 2003): First, are models that include information about the previous trial superior in explaining experimental data compared to models that do not account for this factor? Second, can inter-trial effects be explained by changes in either the generation of voluntary saccades, inhibitory control, or a combination of both?

To answer these questions, we fitted SERIA models that explain actions and RT not only as a function of the current trial type, but also as a function of the previous trial. For this, all trials were divided into four different conditions, according to the cue displayed (pro- or antisaccade) and whether it was a switch or a repeat trial. Although a completely different set of parameters could operate in each condition, this seems biologically implausible and our goal was to identify which parameters could be fixed across conditions, without compromising the ability of the models to parsimoniously explain participants’ behavior. Based on our previous findings (Aponte et al., 2018), we constrained our model space so that the parameters of the early unit, as well as the no-decision time, the probability of an early outlier, and the delay of the late units (Aponte et al., 2017) were fixed across all conditions.

The first model that we considered did not account for the effect of the previous trial. However, we allowed both the inhibitory and the two late units to vary between pro- and antisaccade trials. Thereby, this model included in addition to the constrained parameters (e.g., the 2 parameters for the early unit) 12 parameters for the late and inhibitory units (2×3=6 per trial type). We refer to this model as the *no-switch* model.

Next, we considered the hypothesis that the late units but not the inhibitory unit could change on switch trials. Compared to the *no-switch* model, this model required 2×4=8 additional parameters for the late pro- and antisaccade units on switch and repeat trials. We refer to it as the *switch:late* model. By contrast, in the *switch:inhib*. model we allowed the inhibitory unit but not the late units to differ between switch and repeat trials. This required only 2×2=4 extra parameters compared to the *no-switch* model. Finally, we combined the last two models into the *switch:inhib.+late* model, by permitting the two late units and the inhibitory unit to vary between switch and repeat trials. Hence, this model required (4×2)+(2×2)=12 more parameters than the *no-switch* model.

### Model fitting

All models were estimated using the techniques described in our previous studies (Aponte et al., 2017; 2018). Data from all subjects were entered simultaneously into a hierarchical model presented in Aponte et al. (2018). Samples from the posterior distribution were drawn using the Metropolis-Hasting algorithm. The evidence or marginal likelihood of a model was computed using thermodynamic integration (Gelman and Meng, 1998; Aponte et al., 2016) with 16 parallel chains ordered according to the temperature schedule in Calderhead and Girolami (2009). The algorithm was run for 130000 iterations, from which the last 30000 were used to compute summary statistics. The implementation of the models and inference is available in the open source TAPAS toolbox (http://translationalneuromodeling.org/tapas/).

We were interested in several model-based statistics derived from the fits. First, we evaluated the probability of an inhibition failure, defined as the probability that the early unit hits threshold before all other units:

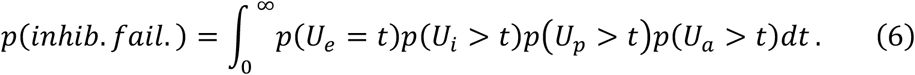

Inhibition failures are fast, reflexive prosaccades, which are correct on prosaccade trials and errors on antisaccade trials.

We also report the conditional probability of a late prosaccade, defined as the probability that the late prosaccade unit hits threshold before the antisaccade unit:

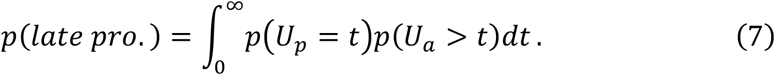

Note that the conditional probability of an antisaccade is defined as

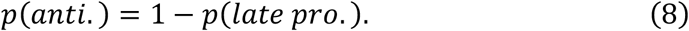

We were also interested in the expected hit times of the late units, defined as

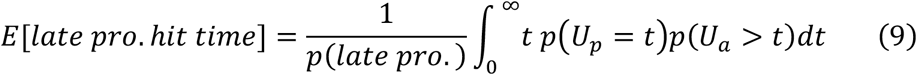

and analogously so for antisaccades. This quantity is the expected hit time of the late prosaccade unit, conditioned on the antisaccade unit arriving at a later point. We report this statistic, as it conveys an interpretable quantity that can be readily compared to experimental data.

## Results

From the 25 participants recruited, two subjects were not included in the final analysis. One subject was excluded because of incomplete data, and the second because in two blocks more than 50% of the trials were either invalid or directly followed an invalid trial.

In the following, we report Task 1 and 2 separately. First, classical statistical analyses of mean RT and ER are presented. These are followed by model-based analyses, in which we compare the *no-switch* and *switch* models using quantitative Bayesian model comparison. We then restrict our attention to *switch* models and explore them in detail, using posterior predictive fits (Gelman et al., 2003) to test when and why individual models fail to predict participants’ behavior.

### Task 1

In Task 1, roughly 4% of the trials were discarded.

#### Error rate and reaction times

Mean RT, ER and switch cost in all conditions are displayed in Fig. 3. On average, participants made significantly more errors on anti- (22 ±21%) than on prosaccade trials (14 ± 14%; X^2^ = (1,276) = 146.2, *p* < 10^-5^), and on switch (27 ± 19%) than on repeat trials (10 ± 13%; X^2^ = (1,276) = 406.6, *p* < 10^-5^). There was a significant interaction between TT and SWITCH (X^2^ = (1,276) = 8.4, *p* = 0.003) demonstrating different switch costs on prosaccade trials compared to antisaccade trials. The antisaccade switch cost (19%) was 4% higher than the prosaccade switch cost (15%).

**Figure 3:**
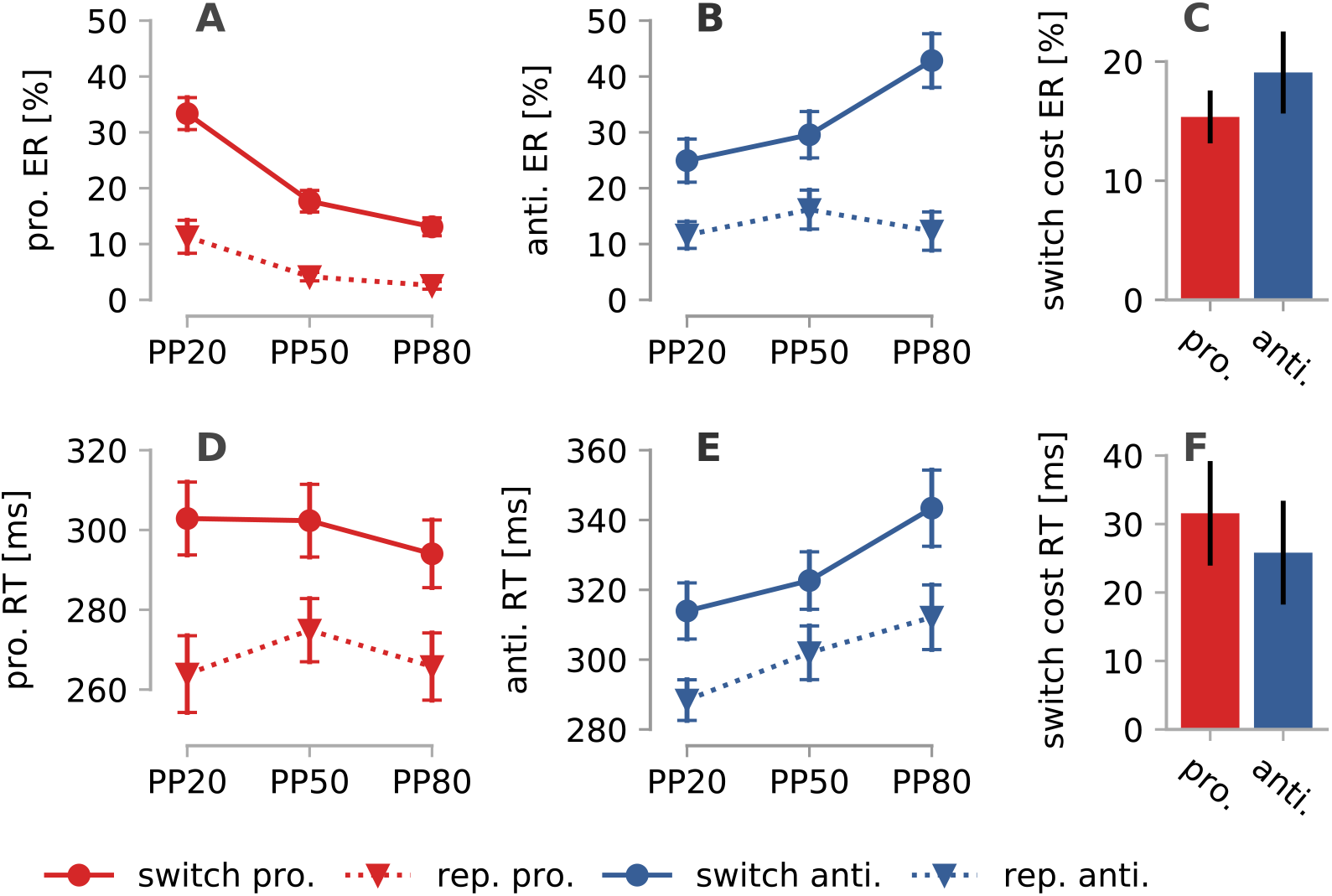
Error rate (ER) and reaction time (RT) in Task 1. **A.** Mean ER on prosaccade trials. **B.** Mean ER on antisaccade trials. **C.** ER switch costs. **D.** Mean RT on prosaccade trials. **E.** Mean RT on antisaccade trials. **F.** RT switch cost. Error bars display the sem. PP: prosaccade trial probability.

Regarding RT, antisaccades (313 ± 44*ms*) were significantly slower than prosaccades (284 ± 45*ms*; *F*_1,242_ = 57.8, *p* < 10^-5^); switch trials (313 ± 46*ms*) were slower than repeat trials (285 ± 43*ms*; *F*_1,242_ = 53.6, *p* < 10^-5^). The interaction between TT and SWITCH was not significant (*F*_1,242_ = 0.5, *p* = 0.463), or, in other words, the antisaccade switch cost (26*ms*) did not significantly differ from the prosaccade switch cost (32*ms*).

#### SERIA – model comparison

All models were initially evaluated according to their log evidence or log marginal likelihood, which corresponds to the accuracy or expected log likelihood of a model adjusted by its complexity (Stephan et al., 2009). Table 1 reports the evidence and accuracy of all models in log units. The model with the highest evidence was the *switch:inhib.+late* model (LME=-16153.3, ΔLME>44 log units compared to all other models), which also obtained the highest accuracy. Note that this model was heavily penalized (accuracy-evidence=940) compared to the simpler models *no-switch* (accuracy-evidence=782) *switch: late* (accuracy-evidence=922) and *switch: inhib*. (accuracy-evidence=834).

**Table 1.** Model comparison. Log model evidence (LME) and expected log-likelihood or accuracy (displayed for comparison) are listed for the four models tested. The highest evidence and accuracy from the switch models are highlighted in bold.

|  | Accuracy | LME |
| --- | --- | --- |
| *no-switch* | -15673 | -16455 |
| *switch:late* | -15231 | -16175 |
| *switch:inhib.* | -15400 | -16235 |
| *switch:inhib.+late* | **-15212** | **-16153** |

The predictive fits of all models are displayed in Fig. 4. These represent the expected predictive distribution estimated from posterior samples. Visual inspection suggests that while the *no-switch* model failed to capture the distribution of switch prosaccades (Fig. 4C), the *switch:inhib*. model failed to capture the distribution of late responses, and particularly so on prosaccade trials (Fig. 4A and C). The *switch:late* model made a better job regarding late saccades, but it did not capture early errors on antisaccade trials (Fig. 4D). Finally, the *switch:late+inhib*. model was able to accommodate most relevant features of subjects’ behavior.

**Figure 4:**
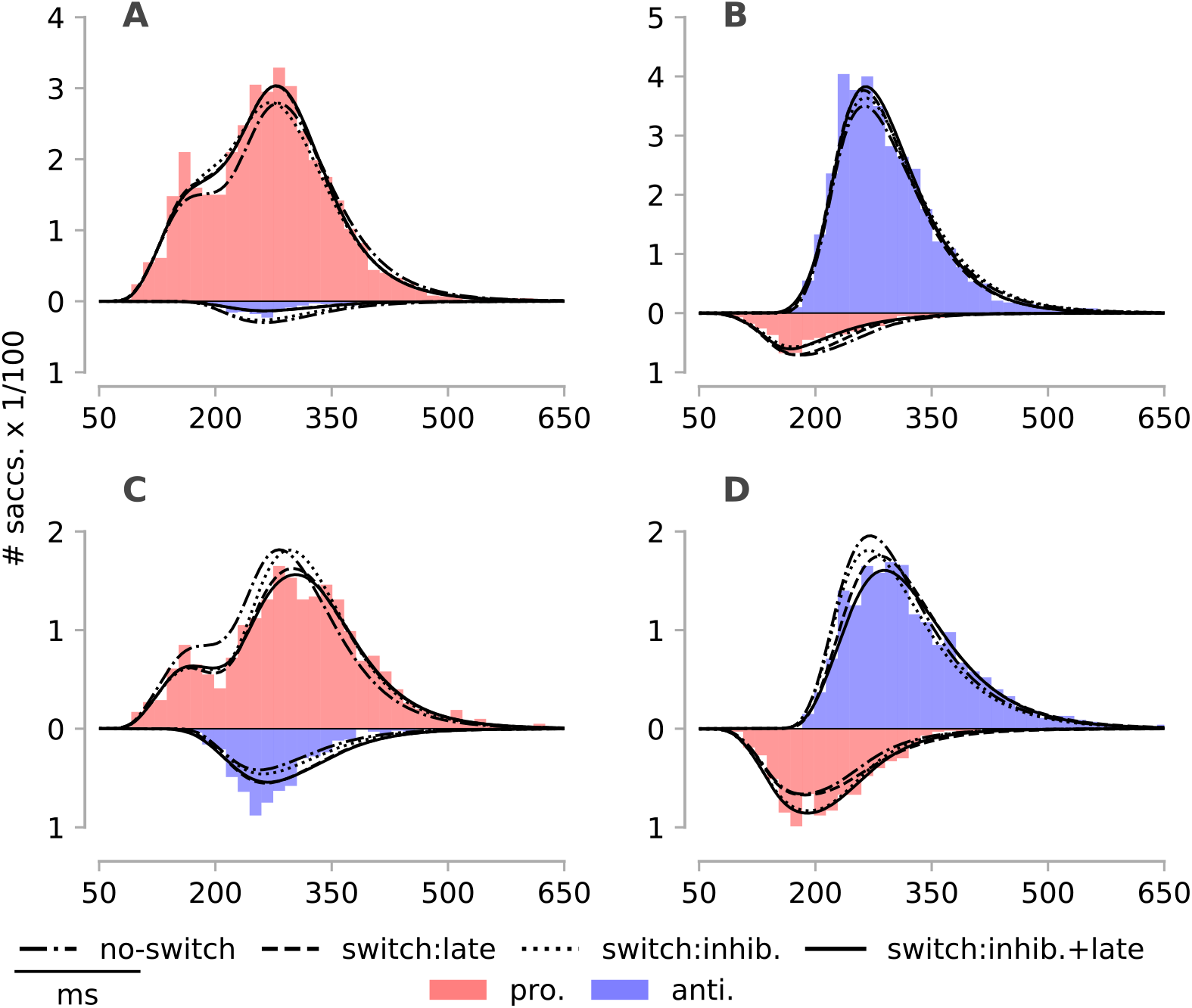
Histogram of the empirical reaction time (RT) and model fits. The empirical RT distribution of prosaccades is displayed in red, and the RT distribution of antisaccades in blue. Errors are displayed in the negative half plane. The weighted posterior predictive distributions of models *no-switch, switch:late, switch:inhib*. and *switch:late+inhib*. are plotted in different line styles. **A.** Prosaccade repeat trials. **B.** Antisaccade repeat trials. **C.** Prosaccade switch trials. **D.** Antisaccade switch trials.

Fig. 5 displays the ER and RT switch costs predicted by all *switch* models. Clearly, only model *switch:inhib.+late* was able to capture switch costs on both pro- and antisaccade trials, whereas model *switch:late* and *switch:inhib*. only correctly explained ER and RT in one of the trial types.

**Figure 5:**
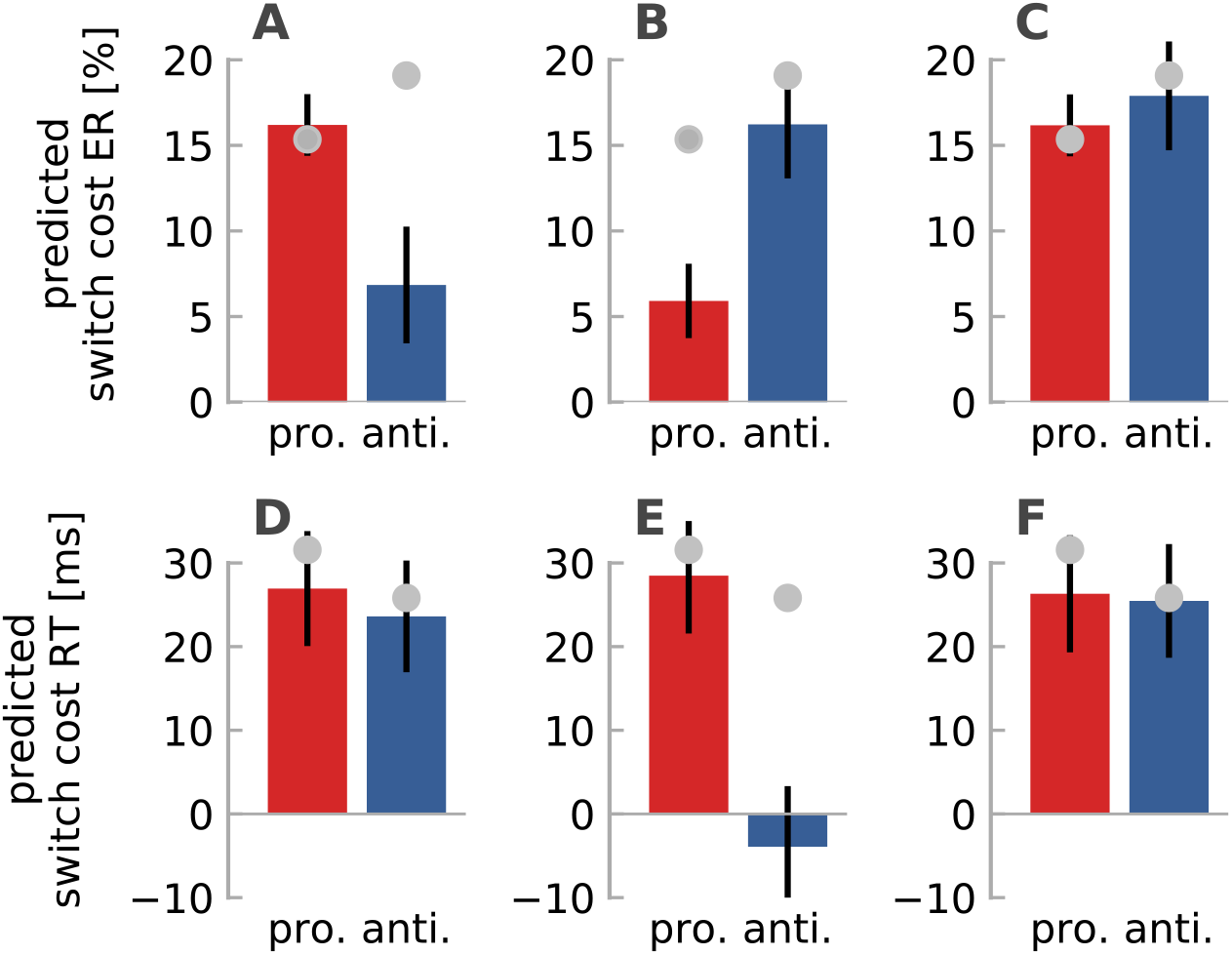
Predicted error rate (ER) and reaction time (RT) switch costs. ER switch cost predicted by the *switch* models. Empirical switch costs (Fig. 3C and 3F) are displayed as gray circles. **A.** *switch:late*. **B.** *switch:inhib.*. **C.** *switch:inhib.+late*. While the *switch:late* model correctly predicted the ER switch costs on prosaccade trials, antisaccade ER costs were clearly underestimated. By contrast, the *switch:inhib*. model captured ER costs on anti- but not on prosaccade trials. The *switch:inhib.+late* made a good job on pro- and antisaccade trials. **D-F.** RT switch cost predicted by the *switch* models. **D.** *switch:late*. **E.** *switch:inhib.*. **F.** *switch:inhib.+late*. The *switch:late* and *switch:inhib.+late* but not the *switch:inhib*. models captured RT switch costs in both pro- and antisaccade trials. Error bars depict the sem. of the model predictions.

#### SERIA – parameter estimates

According to SERIA, there are two types of errors on antisaccade trials: inhibition failures and late prosaccades. To disentangle how these two types of errors contributed to the antisaccade switch cost, we turned first our attention to the probability of an inhibition failure (see Eq. 6), defined as the probability that the early unit hits threshold before all other units. The *switch:inhib.+late* model predicted that, on prosaccade trials, 28±19% of all saccades were inhibition failures, whereas this number was lower on antisaccade trials (21±18%). The effect of switching on pro- (*X*^2^(1,138) = 107.9, *p* < 10^-3^) and antisaccades trials (*X*^2^(1,138) = 229.2, *p* < 10^-5^) was significant. When considered together, we found a significant interaction between TT and SWITCH (*X*^2^(1,276) = 302.1, *p* < 10^-5^). Concretely, prosaccade trials induced more inhibition failures on the next trial, regardless of trial type (pro. switch cost=-18%; anti. switch cost=19%; Fig. 6A).

**Figure 6:**
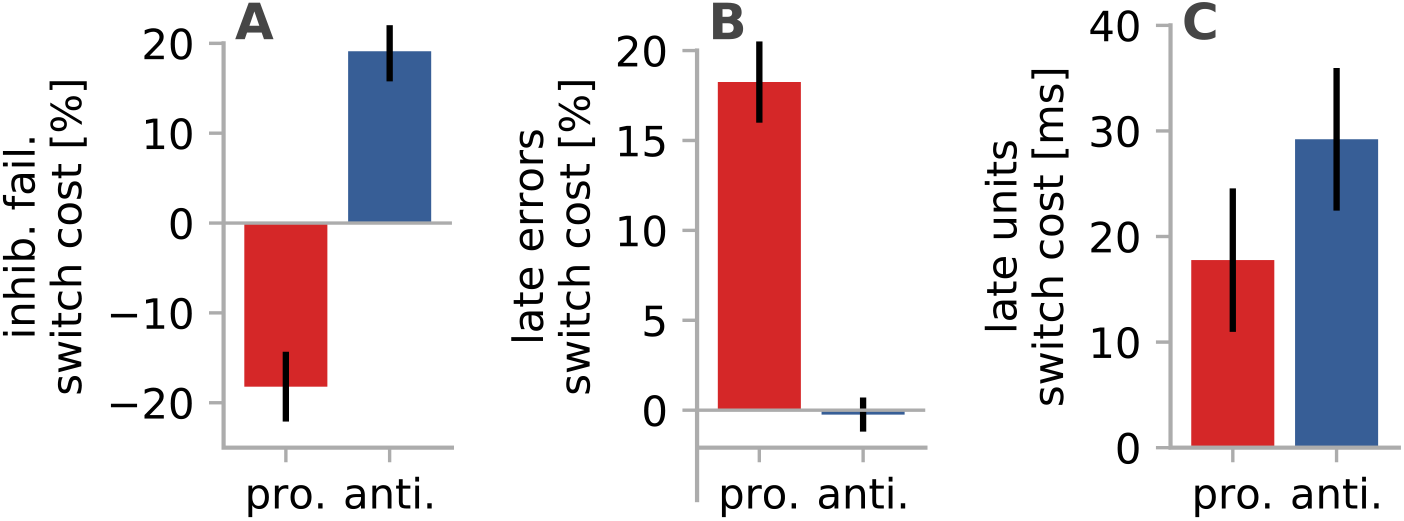
Switch costs in task 1. **A.** Inhibition failures switch cost according to model *m*_4_ *switch:inhib.+late* (Eq. 6). **B.** Late error switch cost (Eq. 7 and 8). **C.** Late units’ hit time (Eq. 9) switch cost. Error bars represent the sem..

This suggested the same number of inhibition failures following a prosaccade trial, regardless of the next trial type. To explore this hypothesis, we fitted a model in which the inhibitory unit was fixed across switch prosaccade and repeat antisaccade trials and across repeat prosaccade and switch antisaccades. The evidence of this post-hoc model was higher than the evidence of *switch:inhib.+late* model (Δ*LME* = 12.1). Qualitatively, there were no large differences in the predictions and parameters of the model. Thus, regardless of trial type, early reactions were similarly inhibited after an antisaccade trial compared to a prosaccade trial.

Next, we submitted the probability of late errors (Fig. 6B; Eq. 7 and 8) on pro- (19±15%) and antisaccade (4±5%) trials to a single GLME. This revealed a significant interaction between SWITCH and TT (*X*^2^(1,276) = 63.0, *p* < 10^-5^). The mean late error switch cost on prosaccade trials was 18%, whereas on antisaccade trials, it was less than 1%. When late ER on antisaccade trials was analyzed separately, the factor SWITCH was not significant (*X*^2^(1,138) = 0.1, *p* = 0.81).

Finally, we investigated the hit time of the late units (Fig. 6C; Eq. 9). Switch late reactions (335±42ms) were significantly (*F*_1,248_ = 81.9, *p* < 10^-5^) slower than repeat reactions (312±36ms). The late prosaccade RT switch cost (18ms) was lower than the antisaccade unit RT cost (29ms) which resulted in a significant interaction between TT and SWITCH (*F*_1,248_ = 4.8, *p* = 0.028).

### Task 2

In Task 2, around 9% of all trials were discarded. At most, 35% of all trials in a single block were excluded.

#### Error rate and reaction time

In this condition (Fig. 7), subjects made significantly fewer errors on pro- (2 ± 4%; Fig. 7A) than on antisaccade trials (13 ± 13%; X^2^ = (1,276) = 297.4, *p* < 10^-5^; 7B), and on repeat (5 ± 10%) than on switch trials (10 ± 12%; X^2^ = (1,276) = 77.4,*p* < 10^-5^). There was a significant interaction between SWITCH and TT (X^2^ = (1,276) = 6.3,*p* = 0.011; Fig. 7C) driven by larger switch costs on antisaccade trials (8%) than on prosaccade trials (3%).

**Figure 7:**
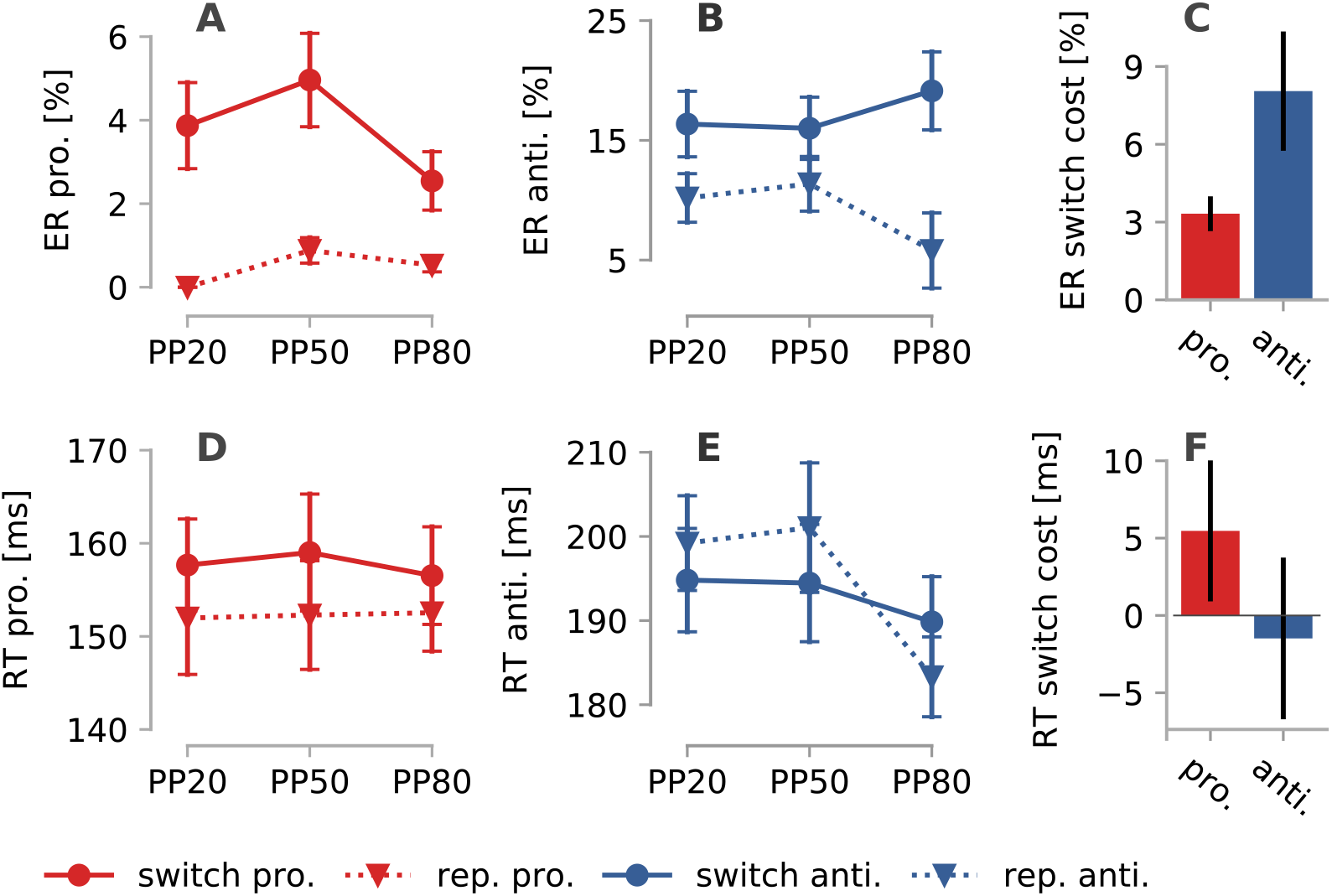
Error rate (ER) and reaction times (RT) in Task 2. **A.** Mean ER on prosaccade trials. **B.** Mean ER on antisaccade trials. **C.** ER switch costs. **D.** Mean RT on prosaccade trials. **E.** Mean RT on antisaccade trials. **E.** RT switch cost. Error bars display the sem. PP: prosaccade probability.

Prosaccades (Fig. 7D; 155 ± 26*ms*) were faster than antisaccades (Fig. 7E; 194 ± 30*ms*; *F*_1,242_ = 385.8, *p* < 10^-5^), but neither the effect of SWITCH (*F*_1,242_ = 1.0, *p* = 0.314) nor the interaction between SWITCH and TT (*F*_1,242_ = 3.0, *p* = 0.079) were significant (Fig. 7F). Nevertheless, we submitted pro- and antisaccades to two separate GLME. As shown in Fig. 7F, prosaccades were significantly faster on repeat than on switch trials (Δ*RT* = 5*ms*; *F*_1,110_ = 6.4,*p* = 0.012), but there was no significant difference on antisaccade trials (Δ*RT* = −1*ms*; *F*_1,110_ = 0.2, *p* = 0.576), although switch antisaccades were slightly faster than repeat antisaccades.

#### SERIA - Model comparison

Contrary to the findings in Task 1, the model with the highest evidence (*no-switch*) not account for any switch cost (Table 2). The second best model was the *switch:inhib*. model, in which the inhibitory unit was allowed to change across all four possible conditions, but the late units could not differ between switch and repeat trials. The difference in LME between *no-switch* and *switch:inhib*. models is explained by a much larger penalty for the latter model (749 and 813 respectively).

**Table 2.**
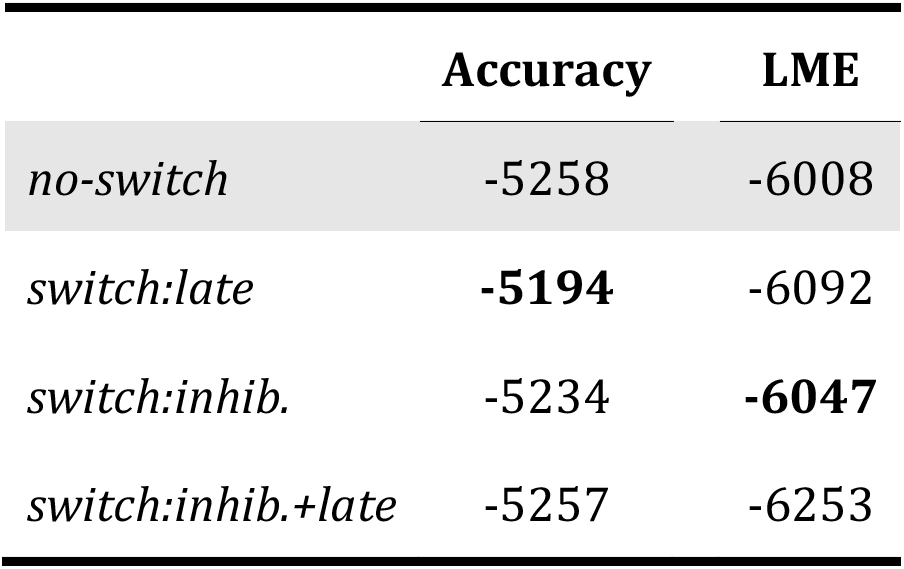
Model comparison. Log model evidence (LME) and expected log-likelihood or accuracy are listed for the four models tested. The highest evidence and accuracy from the switch models are highlighted in bold.

|  | Accuracy | LME |
| --- | --- | --- |
| *no-switch* | -5258 | -6008 |
| *switch:late* | **-5194** | -6092 |
| *switch:inhib.* | -5234 | **-6047** |
| *switch:inhib.+late* | -5257 | -6253 |

All four models fitted RT and ER well in Task 2 (Fig. 8), with no obvious difference between them. This reflects the subtle effects of switching in Task 2. The *switch:late, switch:inhib.+late* predicted switch costs most accurately (Fig. 9A and C), but had a lower evidence than the *switch:inhib*. model.

**Figure 8:**
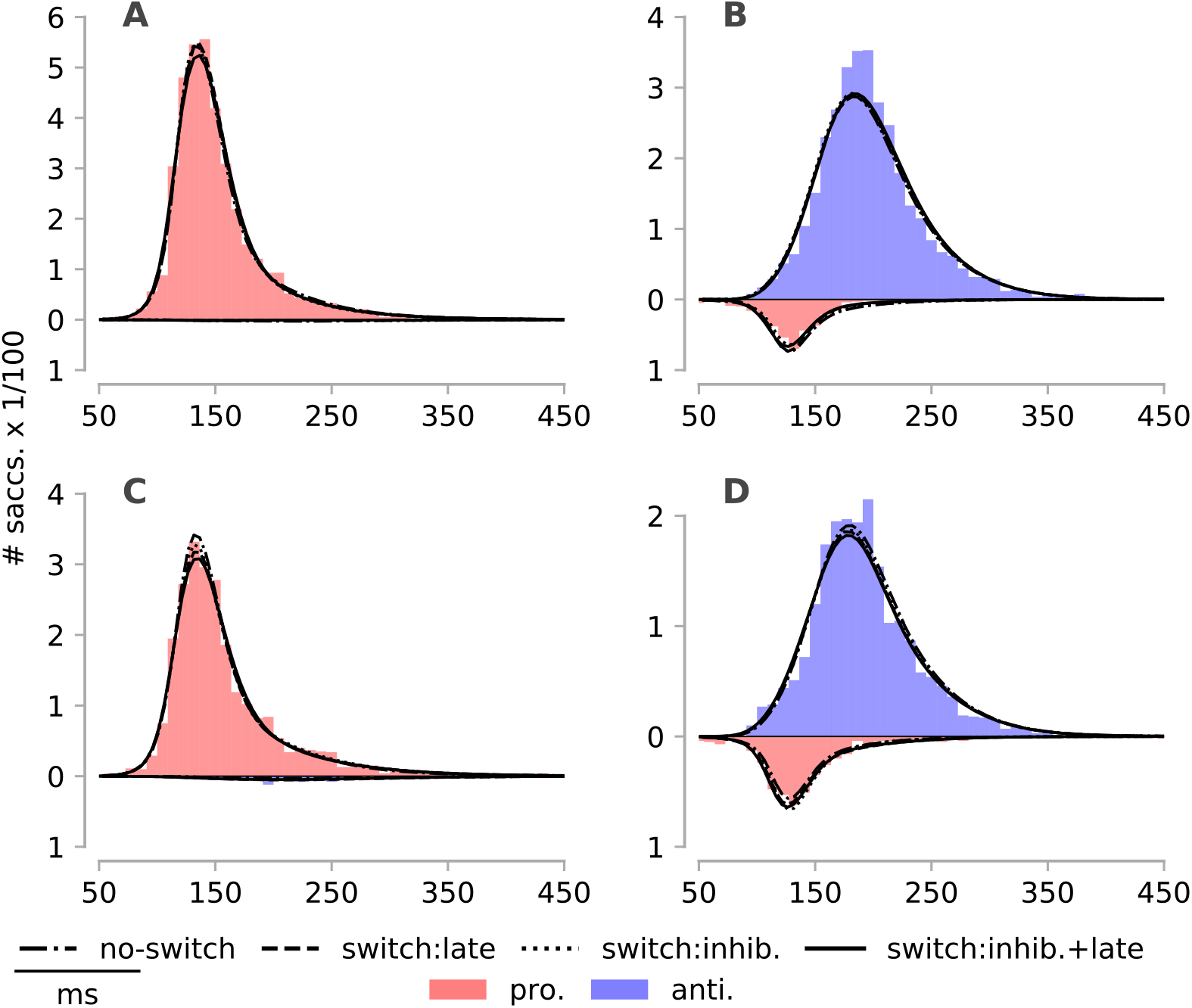
RT histograms and predictive model fits in Task 2. Similar to Fig. 4. **A.** Prosaccade repeat trials. **B.** Antisaccade repeat trials. **C.** Prosaccade switch trials. **D.** Antisaccade switch trials. With the exception of prosaccade switch trials (C), all models generated similar fits.

Because the classical analysis clearly demonstrated the presence of switch costs, we continued to investigate the best *switch* model. Hence, we proceeded to discuss switch costs in Task 2 based on the *switch:inhib*. model. We come back to models *switch:late* and *switch:inhib.+late* below.

**Figure 9:**
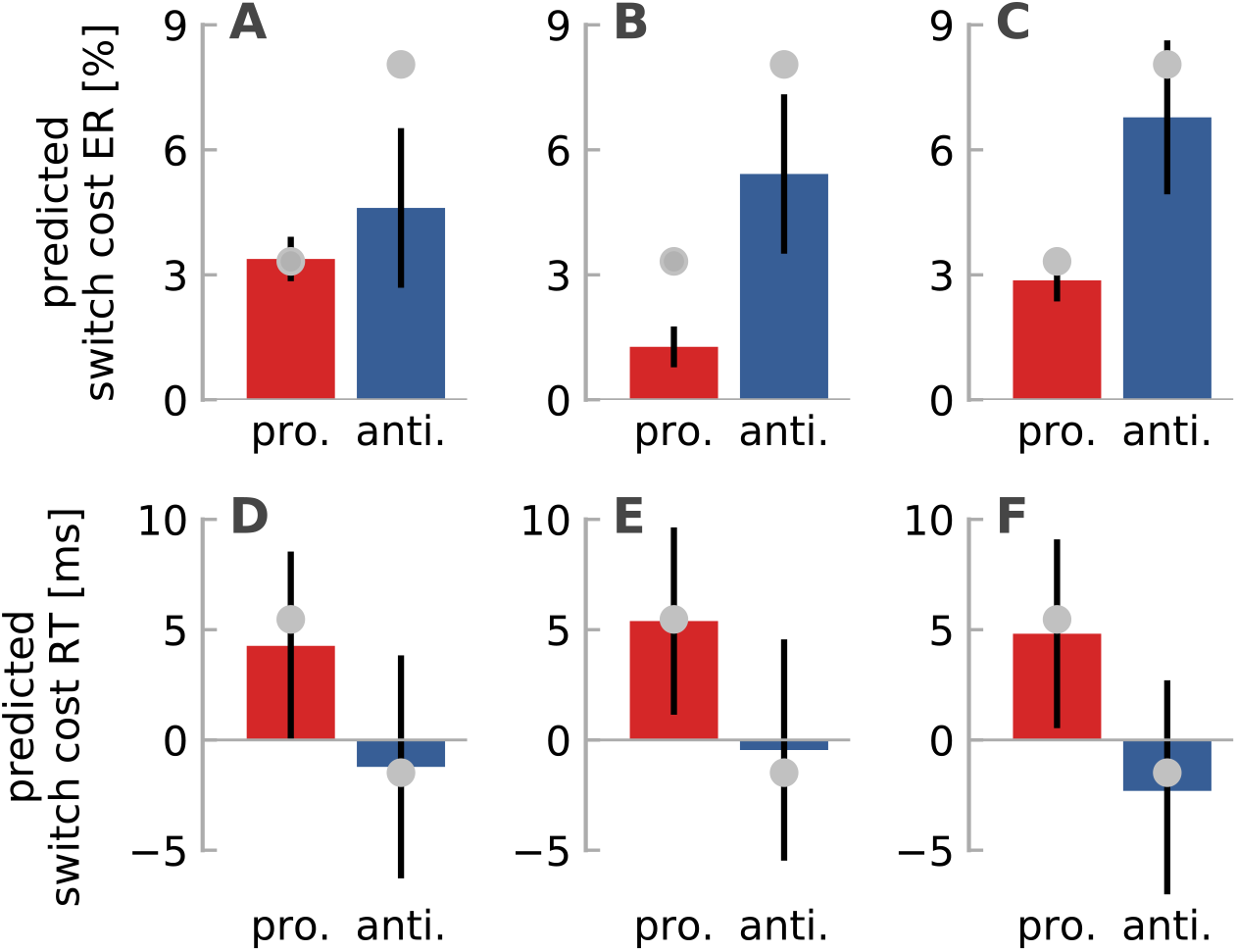
Model predictions. **Top.** Predicted ER switch cost. **A.** *switch:late*. **B.** *switch:inhib.*. **C.** switch:inhib.+late. Bottom. Predicted RT switch cost. **D.** *switch:late*. **E.** *switch:inhib.*. **F.** switch:inhib.+late.

Qualitatively, the *switch:inhib*. model (Fig. 9B) could reproduce our main behavioral findings: switch trials were characterized by higher ER (10±11%) than repeat trials (6±11%; *X*^2^(1,248) = 58.3,*p* < 10^-5^). Although the predicted switch cost was higher on anti- (5.4%) than on prosaccade trials (1.2%), the interaction between SWITCH and TT was not significant (*X*^2^(1,248) = 0.1,*p* = 0.75) contrary to our behavioral analysis. Moreover, the model clearly underestimated the ER switch cost on pro- and antisaccade trials (Fig. 9B), as discussed in the next section. Regarding RT, the model predicted a positive switch cost on prosaccade trials (5ms, *F*_1,112_ = 9.8, *p* = 0.002), as well as a negative but negligible cost on antisaccades trials (*F*_1,112_ = 0.0, *p* = 0.834).

#### SERIA – model parameters

To understand how the *switch:inhib*. model was able to capture switch costs in Task 2 without postulating changes in the late units, we plotted the probability of inhibition failures on switch and repeat trials (Fig. 10A-B). As in Task 1, saccades that followed prosaccade trials were more likely to be inhibition failures, regardless of trial type (Fig. 10C; interaction TT*SWITCH; *X*^2^(1,248) = 47.7, *p* < 10^-5^). This allowed for more late reactions on switch prosaccade trials, and conversely, more early errors on switch antisaccade trials. Because switch prosaccades yielded more late reactions than repeat prosaccades, these trials were accompanied by more slow saccades. In summary, prosaccades led to more inhibition failures on the next trial (regardless of trial type). However, the base line of inhibitory control was different on pro- and antisaccade trials, as subjects made roughly 7 times more inhibition failures on prosaccade trials (60±15%) than on antisaccade trials (8±9%).

**Figure 10:**
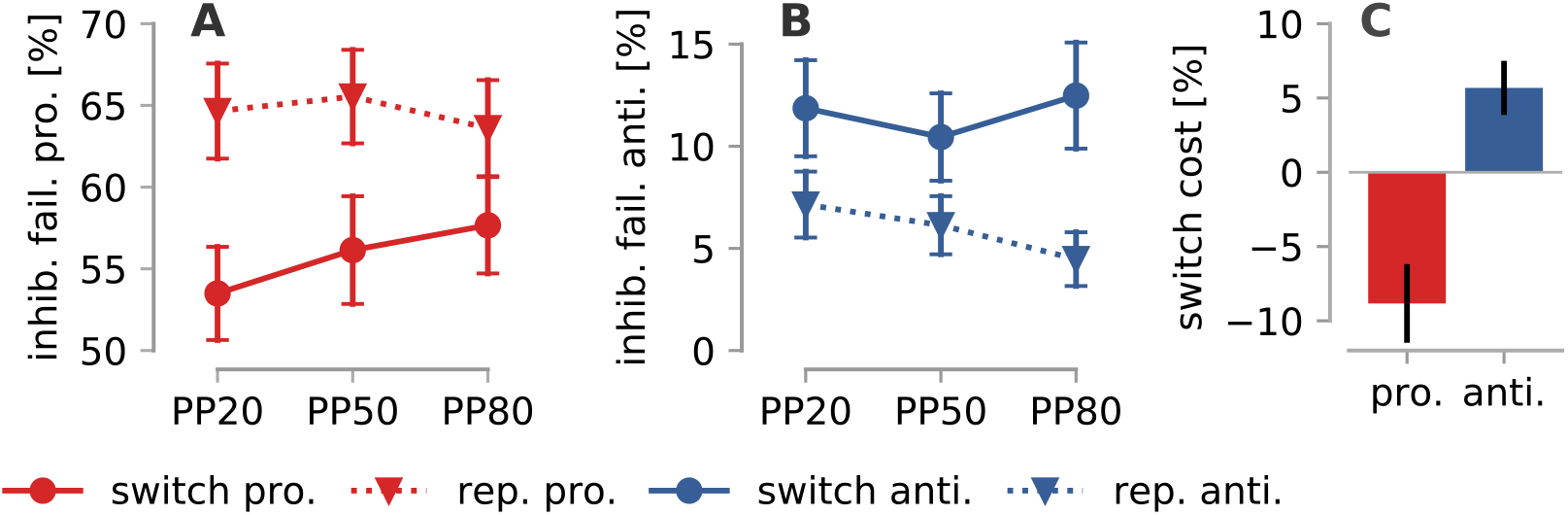
Inhibition failures in Task 2 according to the *switch:inhib*. model. **A.** Predicted probability of an inhibition failure on a prosaccade trial. **B.** Inhibition failures on antisaccade trials. **C.** Inhibition switch cost on pro- (9%) and antisaccade (6%) trials. Error bars display the sem..

#### Prosaccade and antisaccade ER switch cost

As illustrated above (Fig. 9B), the *switch:inhib* model underestimated the ER switch cost and its predictions did not support a significant interaction between the factors SWITCH and TT. Careful examination revealed that although the *switch:inhib*. model could partially account for the ER on prosaccade trials (Fig 11A-B; repeat: 1.6%; switch: 2.9%), it could not fully capture the eightfold increase in ER between prosaccade repeat (0.47%) and switch trials (3.79%).

**Figure 11:**
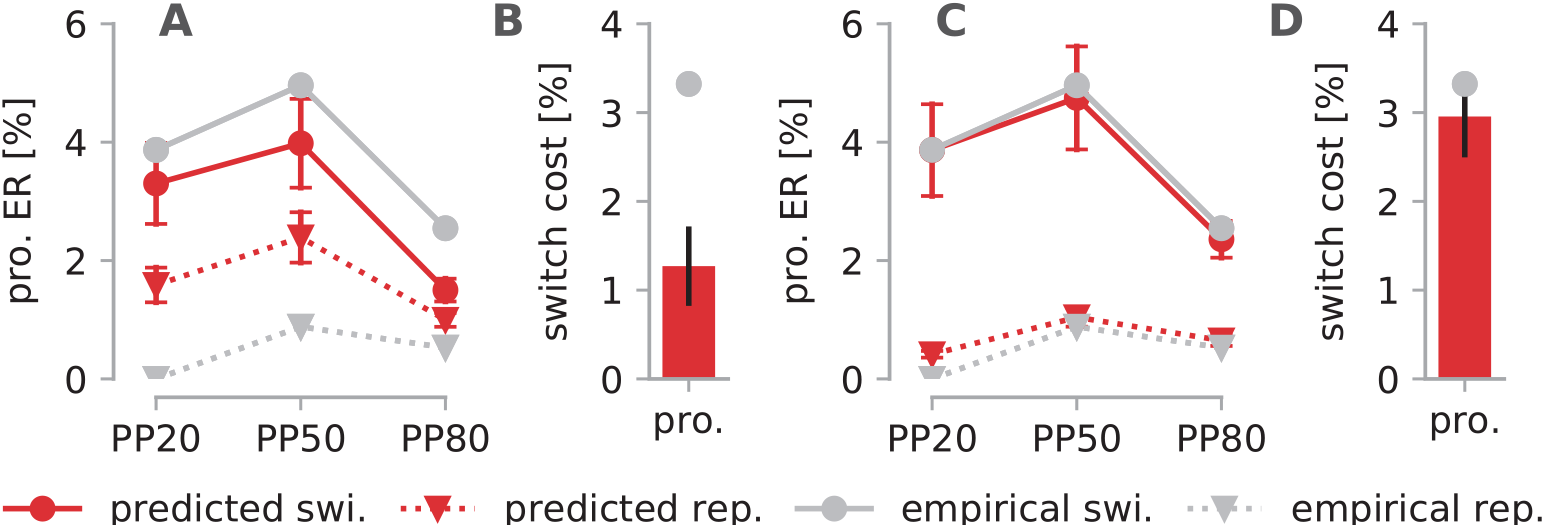
Predicted and empirical ER and switch cost on prosaccade trials. **A. *switch:inhib*. predictions.** The *switch:inhib*. model accounts for the switch cost only through changes in inhibitory control. **B. *switch:inhib*. prosaccade ER switch cost.** Although this model does capture a fraction of the switch cost, it is limited by the proportion of inhibition failures on repeat and switch trials. For visualization the empirical switch cost is displayed as a gray circle. **C. *switch:inhib.+anti*. predictions.** In the *switch:inhib.+anti*. model, the antisaccade unit is allowed to vary between prosaccade switch and repeat trials. In this case, the predicted error rate on repeat trials is closer to the empirical error rate. **D. *switch:inhib.+anti* prosaccade ER switch cost.** Similar to panel B. Error bars display the sem. of the model predictions.

According to SERIA, an error on a prosaccade trial can almost only^1^ be generated when an early response is inhibited and the antisaccade unit hits threshold before the late prosaccade unit. Thereby, the prosaccade ER is *approximately* equal to

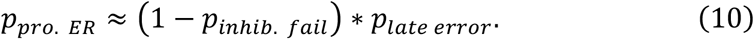

In the *switch:inhib* model, the late units are assumed to not change across switch and repeat trials and thereby, an eightfold increase in the ER is only possible if there is an eightfold change in the probability of a late action (i.e., 1 − *P_inhib. fail_*; Eq. 10). However, such a large change is not possible given the predicted number of inhibition failures on prosaccade trials (60%; see Fig 10A). Thus, higher ER on switch trials can only be explained by changes in the late units.

To account for this cost, we considered a model (*switch:inhib.+anti.*) in which we allowed the parameters of the antisaccade unit to vary between switch and repeat prosaccade trials. These parameters control the probability and RT of errors on prosaccade trials but have no influence on antisaccade trials. As displayed in Fig. 11C-D, the predicted ER on switch and repeat trials using the *switch:inhib.+anti*. model was 3.67% and 0.67%, respectively. When we considered again the interaction between the factors SWITCH and TT using the predicted ER of the *switch:inhib.+anti*. model, this was significant (*X*^2^(1,276) = 20.5,*p* < 10^-5^). Nevertheless, the *switch:inhib.+anti*. model had a lower LME than the *switch:inhib*. model (ΔLME=67.0).

Regarding antisaccade trials, the ER switch cost was underestimated by the *switch:inhib*. model (empirical 8.1%, predicted 5.3%). However, as shown in Fig. 12, this was mainly due to the PP80 condition, in which the empirical ER in repeat trials was lower than predicted by the model. Note that this condition is by design much less frequent than the others, and thereby the empirical mean ER displays high uncertainty. Taken together, our analyses demonstrate that, similarly to Task 1, alternating from an antisaccade to a voluntary prosaccade induces more *late errors* compared to repeat prosaccades. However, there is no *late error cost* associated with alternating from a prosaccade to an antisaccade trial.

**Figure 12:**
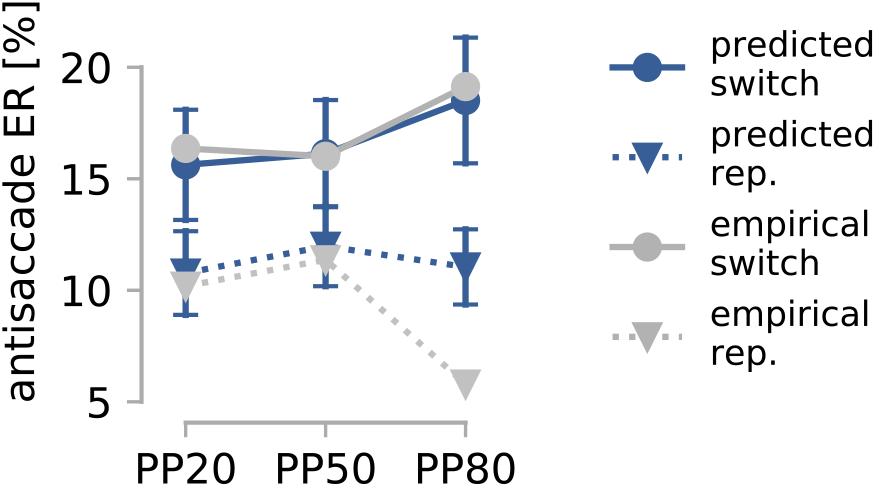
Predicted and empirical ER on antisaccade trials. Predictions were obtained using the *switch:inhib*. model, in which the late units are assumed to not change across switch and repeat trials. The model overestimated the empirical ER for repeat trials in the PP80 condition only. Error bars display the sem. of the model predictions.

## Discussion

Here, we investigated switch costs in the mixed antisaccade task with the help of a computational model. This allowed us to accurately quantify to what extent task switching affects the inhibition of habitual responses (early prosaccades) and voluntary behavior (late pro- and antisaccades). Modeling revealed two main distinguishable effects: First, in Task 1 but not in Task 2, switch trials engendered RT costs in late, voluntary saccades. Second, in both tasks, early reactions that followed prosaccade trials were less likely to be inhibited compared to saccades that followed antisaccade trials. Can SERIA accommodate all or some of the predictions of the task-set reconfiguration and the task-inertia hypotheses? Does SERIA provide an alternative or more fined-grained explanation for these predictions? In the following, we discuss the answer to these questions.

### Switch costs in the antisaccade task

Findings in the mixed antisaccade task can be divided into to two main groups. Early studies (e.g., Barton et al., 2002; Cherkasova et al., 2002; Manoach et al., 2002; Fecteau et al., 2004) reported positive prosaccade RT switch costs, negative antisaccade RT costs, as well as higher ER in switch trials regardless of trial type. More recently, Heath and Weiler (e.g., Weiler and Heath, 2012a; Weiler et al., 2015) reported positive RT switch costs on prosaccade trials, and no RT switch costs on antisaccade trials. Again, all switch trials were characterized by higher ER.

Our empirical findings are well in line with these previous reports. Regarding Task 1, positive switch costs in pro- and antisaccade trials have been previously demonstrated in a similar design by Barton et al. (2006a); see also (Hunt and Klein, 2002), who displayed the task cue 200ms in advance of the peripheral stimulus. In Task 2, we found non-significant negative antisaccade RT switch costs, as well as significant positive prosaccade RT switch costs. This is congruent with the unidirectional switch costs reported by Weiler and Heath (2012a).

Based on SERIA, we proposed three models or hypotheses to explain these findings: (i) the *switch:inhib*. model in which only the parameters of the inhibitory unit could change across switch and repeat trials; (ii) the *switch:late* model in which the late units but not the inhibitory unit were allowed to vary across conditions; and (iii), the *switch:inhib.+late* model which combines both hypotheses.

Quantitative Bayesian model selection and qualitative posterior predictive checks (Gelman et al., 2003; Gelman and Shalizi, 2013) indicated that in Task 1 the *switch:inhib.+late* model accounted best for participants’ ER and RT. In Task 2, the model with the highest evidence did not allow for any switch cost. However, in the *switch* family, the *switch:inhib*. model obtained the highest evidence. Qualitatively, this model could fit RT switch costs in pro- and antisaccade trials, and, after an extension, it could fit prosaccade ER switch costs.

SERIA demonstrates therefore that there is a cost associated with switching between voluntary pro- and antisaccades, and that this cost is only observable in Task 1. In particular, we could show that in this task not only switch antisaccades had a higher latency than repeat antisaccades, but *late* switch prosaccades were also delayed compared to repeat prosaccades. In Task 2, SERIA accounted for pro- and antisaccade switch costs without postulating any change in the late units. Fundamentally, this is compatible with the main prediction of the task-set reconfiguration hypothesis (Rogers and Monsell, 1995; Meiran, 1996), which postulates that switching between task-sets is time consuming, but can be done in advance of the response cue.

In addition to the switch cost associated with voluntary actions, we found that there was a consistent inter-trial effect on inhibitory control in Task 1 and 2. Specifically, we found that inhibition failures were more likely after prosaccade trials than after antisaccade trials, regardless of the current trial type. This observation is a direct prediction of the task inertia hypothesis according to which “switch costs should change as a function of the task that participants are switching from, not as a function of the task they are switching to” (Wylie and Allport, 2000). A second prediction of this hypothesis that is confirmed by our modeling is that inter-trial effects persist regardless of the delay between the task-cue and the imperative stimulus (i.e., the peripheral target; Wylie and Allport, 2000).

The answer to our first question (can SERIA accommodate the predictions of the task inertia and task-set reconfiguration hypotheses?) is therefore positive. As in other multiple-component models (reviewed in Schmitz and Voss, 2014), the mechanisms that explains both predictions are assigned to different components: On one hand, asymmetric switch costs that persist regardless of the delay between task and peripheral cues are explained by carry-over inhibition of habitual reactions. On the other hand, higher RT on switch trials in Task 1 (in which subjects cannot prepare their action in advance of the peripheral cue) are assigned to the generation of voluntary actions.

The answer to our second question (how does SERIA explain these predictions?) is more nuanced. Although our modeling is compatible with the predictions of the task inertia hypothesis, SERIA postulates a different mechanism for these inter-trial, carry-over effects. Rather than passive interference between task-set rules (Weiler et al., 2015), our results indicate that the strong inhibition associated with an antisaccade trial reduces the probability of an inhibition failure on the next trial. We come back to this point later.

The mechanism described by SERIA differs also from the theory proposed by Barton et al. (2006a), according to which switch costs are (partially) due to the generalized suppression of the response-system that “affects both the upcoming pro- and antisaccades”. This account is problematic, because generalized inhibition predicts the same effect on switch pro- and antisaccades, keeping their ratio constant compared to repeat trials. By contrasts, in SERIA, stronger inhibition leads to more late responses in prosaccade switch trials, as well as fewer inhibition failures on repeat antisaccade trials, while allowing for negligible antisaccade RT switch costs.

Our results also shed light on the observation that response inhibition in the go/no-go task induce similar RT costs on go trials as antisaccade trials on prosaccade trials (Barton et al., 2006b). In particular, SERIA postulates that the inhibition of early responses in the antisaccade task relies on the same functional mechanism as correct no-go trials in the go/no-go task. Hence, it is a natural prediction that similar carry-over effects should be observed in both paradigms (Barton et al., 2006b).

So far, we have not discussed the negative or paradoxical antisaccade RT switch costs initially reported by Cherkasova et al. (2002). Negative switch costs occur only when the task cue is presented in advance of the peripheral cue (Hunt and Klein, 2002; Barton et al., 2006a), which suggests that negative switch costs are not caused by changes in voluntary action generation. In Supp. Material 1, we demonstrate that the *switch:inhib*. model can simulate negative switch costs, even in the absence of changes in the late units across repeat and switch trials. This is possible because of the non-linear interactions between the antisaccade, the early and the inhibitory units which allow for faster antisaccades in low inhibition conditions (switch trials) compared to high inhibition conditions (repeat trials).

The mechanisms described by SERIA are therefore sufficient to explain the plurality of behavioral findings reported in the antisaccade task: positive switch costs in pro- and antisaccade trials when the task cue is presented shortly before or simultaneously to the peripheral stimulus; and unidirectional switch costs, as well as paradoxical switch costs, when the task cue is presented ahead of the peripheral cue. Next, we discuss in more detail how the *switch:inhib*. model allows for asymmetric switch costs in the absence of changes in the late units.

### Asymmetric costs in habitual and non-habitual responses

A key observation in the task switching literature is that switching from a habitual to a non-habitual response engenders higher costs than switching from a non-habitual to a habitual response (Allport et al., 1994; Wylie and Allport, 2000). SERIA provides a simple mathematical explanation for this phenomenon. The expected RT of dominant or habitual responses can be *approximated* as the mixture of the expected RT of early and late responses

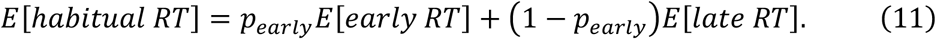

The expected RT of non-habitual responses is given by

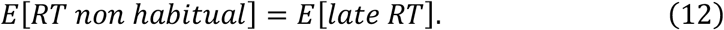

Accordingly, in a transition from a non-habitual to a habitual response, the probability of a late response increases, elevating the overall mean RT, even in the absence of late action switch costs. In the case of a transition from a habitual to a non-habitual response, the RT of non-habitual responses should be equal to the RT of repeat trials. This is how the *switch:inhib*. model explains the positive RT switch cost on prosaccade trials as well as the absence of a significant RT switch cost on antisaccade trials in Task 2. Note that this approximation is invalid in certain circumstances, as demonstrated in Supp. Material 1, in which we show that the *switch:inhib*. model can generate negative antisaccade RT switch costs.

To our knowledge, no other computational model has been used to investigate the inhibition of habitual responses as a component of task switching, nor has this mechanism been used to explain asymmetric task switch costs. Arguably, the reason is that most paradigms used in the task-switching literature do not require actions for which a habitual (or dominant) response is associated, an important exception being the modified Stroop paradigm used originally by (Allport et al., 1994). Nevertheless, it is likely that habitual action inhibition plays an important role in experimental paradigms in which dominant and non-dominant responses are required from participants.

An important qualification here is that while the concept of ‘inhibition’ plays a significant role in the task switching literature (reviewed in Koch et al., 2010), this is usually understood as the ‘proactive interference resulting from having performed a competing task’ (Koch et al., 2010). In the present context, we have used ‘inhibition of habitual responses’ in the narrow sense of ‘motor inhibition’ entailed by the race model proposed by Logan et al. (1984; see Schall et al., 2017). Specifically, inhibition (in this narrow sense) only affects early reactions through the binary, probabilistic competition between independent accumulators.

In summary, switch costs in the mixed antisaccade task can be partially explained by a mechanism that fulfills one of the main predictions of the task inertia hypothesis. However, this mechanism affects only the inhibition of habitual responses, which modulates ER and RT by altering the ratio between habitual and voluntary actions. This form of inhibition should not be confused with proactive interference of tasks-sets, proposed by other theories of task switching.

### Conclusion

Our modeling illustrates how conceptual theories of switch costs can profit from a rigorous formulation in computational terms, as seemingly contradictory hypotheses and findings can be formally unified under a more general theory. In particular, our analysis indicates that alternating between voluntary actions engenders task-set reconfiguration costs, whereas carryover inhibition of habitual responses can explain asymmetric switch costs.

## Acknowledgements

This work was supported by the René and Susanne Braginsky Foundation (KES) and the University of Zurich (KES).

## Supplementary materials

### Supplementary Material 1. Can SERIA explain paradoxical switch costs?

In Task 2, we found a negative but not significant RT switch cost in antisaccade trials, that is, antisaccades that followed prosaccades were faster than repeated antisaccades. Negative antisaccade RT switch costs, called *paradoxical switch costs* (Cherkasova et al., 2002), have been reported in designs in which the trial type is displayed much in advance of the visual cue and are accompanied by an increase in ER. Could the SERIA model account for this finding at all? In order to answer this theoretical question, we set up a simulation (Fig. S1) in which the early prosaccade unit and the late units behaved identically between repeat and switch trials, but the inhibitory unit was allowed to change across both conditions. Although it might be possible to explain antisaccade switch costs relying on the late units, the results from Task 1 and 2 suggest that switching between trial types engender positive but not negative costs on the late units.

Initially (Fig. S1A), we simulated switch and repeat RT distributions varying only the parameters of the inhibitory unit. The RT of switch and repeat antisaccades were 304 and 293ms, whereas the ER were 5 and 28% respectively. The mean antisaccade RT decreased as high latency antisaccades competed with non-inhibited prosaccades. A critical property of this simulation is that the early unit has a sluggish hit time distribution that explains the relatively low error rate (5-28%). Under this condition, when inhibitory control is released, the antisaccade RT distribution is shifted toward lower latencies. As displayed in Fig. S1.B, negative RT switch costs (−15ms) are still possible when the early unit has a narrow distribution, but the ER (rep. 20%, switch 58%) would be much larger than usually reported.

**Figure S1:**
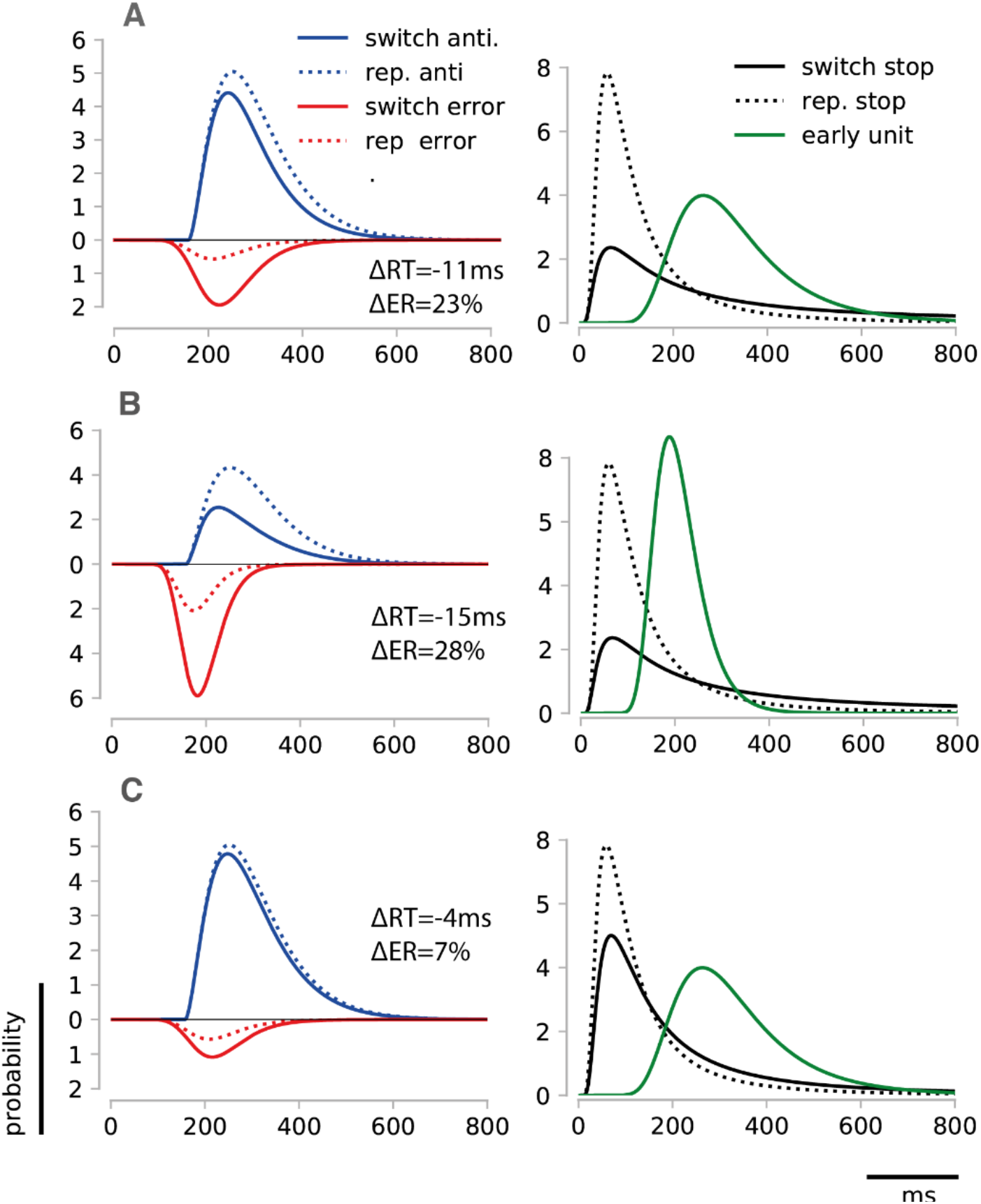
Simulated antisaccade costs. **A. Left:** Simulated antisaccade RT distribution in switch (solid line) and repeat (broken line) trials. Errors are displayed in the bottom half-plane. Probabilities have been scaled by 1000. The RT switch cost was −11ms, and the ER switch cost was 23%. **Right:** Distribution of the hit times of the early and stop unit in repeat and switch trials. Switch antisaccades are characterized by lower inhibitory control. The early unit has a wide distribution that allows for low error rate and negative RT switch cost **B. Left:** Simulated antisaccade RT distribution when the early unit has a narrow distribution. In the switch condition, the ER was 58% and in repeat trials, it was 20%. The RT switch cost was −15ms. **Right.** Distribution of the hit times of the early and stop unit in repeat and switch trials. **C. Left:** Simulated switch cost with moderated release of inhibition in switch trials. RT switch cost: −4ms; ER switch cost: 7%. **Right.** Inhibitory and early units.

What happens when there is a less pronounced release of inhibition in antisaccade switch trials? In Fig. S1C, the latency of the inhibitory unit in switch trials was shorter compared to the first simulation. This reduced the ER in switch trials to 16% (switch cost=7%) but also reduced the negative antisaccade switch cost to only 4ms, demonstrating that for moderate differences in inhibitory control, the paradoxical switch cost is much lower. This potentially accounts for the unidirectional switch cost (Weiler and Heath, 2012a), that is, when there is only a moderate change in inhibitory control, there is no strong change in the RT latency of antisaccades across switch and repeat trials.

## Footnotes

1 It is possible, although highly unlikely, that the antisaccade unit hits threshold before all three other units.

